# Analysis of EEG networks and their correlation with cognitive impairment in preschool children with epilepsy

**DOI:** 10.1101/237172

**Authors:** Eli Kinney-Lang, Michael Yoong, Matthew Hunter, Krishnaraya Kamath Tallur, Jay Shetty, Ailsa McLellan, Richard FM Chin, Javier Escudero

**Affiliations:** School of Engineering, Institute for Digital Communications, The University of Edinburgh, Edinburgh EH9 3FB, United Kingdom; The Muir Maxwell Epilepsy Centre, The University of Edinburgh, Edinburgh EH8 9XD, United Kingdom; Royal Hospital for Sick Children, Edinburgh EH9 1LF, United Kingdom

**Author notes:** Corresponding author Email address (Eli Kinney-Lang). Authors contributed equally to the work (Richard FM Chin, Javier Escudero). Preprint submitted to Epilepsy and Behaviour.

**Keywords:** Network analysis, signal processing, EEG graph networks, paediatric epilepsy, developmental impairment

## Abstract

**Objective:** Cognitive impairment (CI) is common in children with epilepsy and can have devastating effects on their quality of life and that of their family. Early identification of CI is a priority to improve outcomes, but the current gold standard of detection with psychometric assessment is resource intensive and not always available. This paper proposes a novel technique of network analysis using routine clinical electroencephalography (EEG) to help identify CI in children with early-onset epilepsy (CWEOE) (0-5 y.o.).

**Methods:** We analyzed functional networks from routinely acquired EEGs of 51 newly diagnosed CWEOE from a prospective population-based study. Combinations of connectivity metrics (e.g. phase-slope index (PSI)) with sub-network analysis (e.g. cluster-span threshold (CST)) identified significant correlations between network properties and cognition scores via rank correlation analysis with Kendall’s *τ*. Predictive properties were investigated using a 5-fold cross-validated *K*-Nearest Neighbor classification model with normal cognition, mild/moderate CI and severe CI classes.

**Results:** Phase-dependent connectivity metrics had higher sensitivity to cognition scores, with sub-networks identifying significant functional network changes over a broad range of spectral frequencies. Approximately 70.5% of all children were appropriately classified as normal cognition, mild/moderate CI or severe CI using CST network features. CST classification predicted CI classes 55% better than chance, and reduced misclassification penalties by half.

**Conclusions:** CI in CWEOE can be detected with sensitivity at 85% (with respect to identifying either mild/moderate or severe CI) and specificity of 84%, by EEG network analysis.

**Significance:** This study outlines a data-driven methodology for identifying candidate biomarkers of CI in CWEOE from network features. Following additional replication, the proposed method and its use of routinely acquired EEG forms an attractive proposition for supporting clinical assessment of CI.

## Highlights

- EEG network analysis correlates with CI in preschool children with epilepsy
- Classification reveals network features’ predictive potential for CI identification
- Sensitivity to CI improves with dense networks and phase-based connectivity measures

## 1. Introduction

Epilepsy is a complex disease that can have devastating effects on quality of life [1]. Cognitive impairment (CI), which frequently and severely affects quality of life of children and their families, coexists in more than half of children with epilepsy [2, 3, 4, 5]. Timely identification of CI, particularly in children with early-onset epilepsy (CWEOE; epilepsy onset < 5 years of age) is critical because early-life interventions are likely to be more effective, it is the period in which childhood epilepsy is most common, and the most severe forms occur during this time [6, 7, 8]. An estimated 40% of CWEOE have CI [5]. The urgent need for emphasis on early recognition, novel interventions and improved public health strategies for primary and secondary prevention for CI in epilepsy is highlighted in calls to action by august bodies including the International League Against Epilepsy, The Institute of Medicine, and the World Health Organization [9, 10]. Therefore, there is a need to understand the causes of CI and find reliable, affordable and non-invasive markers beyond current standard approaches.

Identification of CI is especially difficult in CWEOE because the gold standard of diagnosis by psychological assessments may not be readily available [11], it is resource intensive, and can be clinically challenging (e.g. introducing potential bias from repeated testing) [11]. Thus, reliable, affordable and rapid CI screening techniques in clinical care are sought after. Such techniques would help focus further medical investigations and resources onto a smaller subgroup, producing efficiency gains and cost savings. Graph network analysis of standard routine clinical EEG recordings is one such potential technique.

Analysis of functional EEG networks offers a data-driven methodology for understanding diverse brain conditions through the lens of network (connectivity) properties [12, 13]. Functional networks examined as graphs are well-established, and provide advantages in understanding changes in connectivity across the brain, e.g. through exploiting properties like small-world topology, connected hubs and modularity [13]. Insights into epilepsy, including the severity of cognitive disturbances, outcomes of epilepsy surgery, and disease duration have been found to correlate with the extent of changes in these functional networks [14]. Recent work has also found network abnormalities can appear in both ictal and interictal states [14]. This supports that network can be distinguished in resting-state EEG [14]. Therefore, functional graph analysis is well positioned as a potential tool to reveal insights into CI in CWEOE.

The aim of this study was to identify a reliable EEG network marker which could help effectively screen for CI in CWEOE. Our hypothesis was two-fold. First, informative network abnormalities could be revealed in CWEOE using graph network analysis on routine clinical EEGs. Second, identified abnormalities could be integrated into a simple machine learning paradigm to demonstrate predictive capabilities with respect to CI. We aimed to utilize a data-driven, quantitative approach to identify potential network markers. Then, we could integrate their information into a simple classification pipeline, which could be readily implemented to support clinical decisions regarding CI. By investigating only routine EEG recordings, we hoped to demonstrate that minimal potential cost and effort would be required to adopt our proposed technique in a clinical setting.

## 2. Methods

The data processing pipeline for each child is summarized in Figure 1.

**Figure 1:**
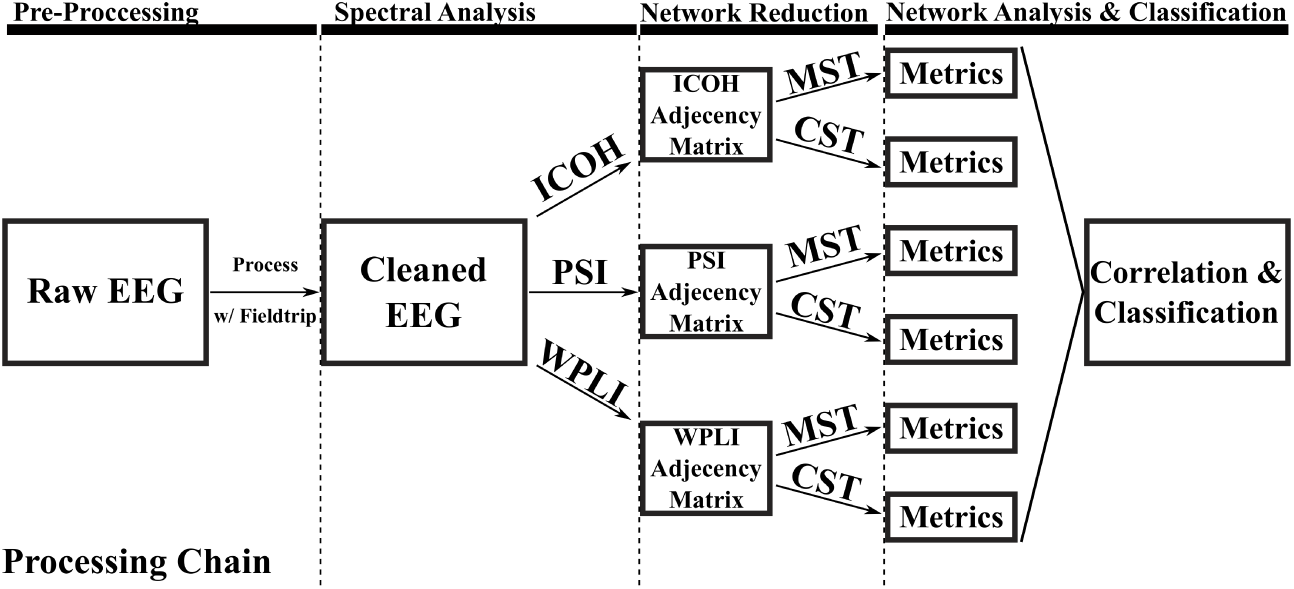
Flowchart of data processing chain for an individual child. ICOH = Imaginary part of coherency, PSI = Phase-slope index, WPLI = Weighted phase-lag index, MST = Minimum Spanning Tree, CST = Cluster-Span Threshold

### 2.1. Dataset

The details on study recruitment and assessments are reported elsewhere [15]. In summary, newly diagnosed CWEOE of mixed epilepsy types and aetiologies were recruited as part of a prospective population-based study of neuro-development in CWEOE. Parents gave approval for use of the standard, resting-state, awake 10-20 EEG their child had as part of their routine clinical care. If a child had multiple EEGs, only the first EEG was used to avoid biasing results toward children with multiple recordings. Additionally, it allowed similar selection of resting-state recordings across all children, e.g. awake resting-state. As such, no EEG recordings of sleep were analysed in this work. All analyses were blinded to any treatment or seizure frequency information. Participants underwent cognitive assessment with age-appropriate standardized tools, e.g. Bayley Scales of Infant and Toddler Development-Third Edition (Bayley-III) and Wechsler Preschool and Primary Scale of Intelligence-Third Edition (WPPSI-III). Children who scored within ±1 standard deviation (SD) of the normative mean were defined as normal, –1 to –2 SD as having mild/moderate CI, and < –2 SD as having severe CI. The cognition scores from Bayley-III and WPPSI-III tests were converted into a normalized standard score measure. Clinical details were collected by members of the research team using a standardized proforma by direct interview of care-givers, medical records and, where possible, patients themselves when they attended for clinical and/or research study assessment.

Table 1 provides the demographic and clinical features for the CWEOE which were included in this study. Given the broad anti-epileptic drug (AED) therapies and aetiologies present in Table 1, potential interactions from AED load or specific aetiology were examined with respect to the designated CI classes (e.g. normal, mild/moderate, severe CI). Using a non-parametric version of the two-way ANOVA (Friedman’s test [16]) on data from Table 1, we revealed no significant interactions between any AED load or specific aetiology with respect to any CI classes. This in turn suggests that the results identified via network analysis are likely driven mainly by cognitive phenomena, as opposed to epileptic syndrome or AED load effects.

**Table 1:** Demographic and clinical feature information of patients, grouped by CI classes of normal, mild/moderate CI, and severe CI. Significant differences between groups with respect to age are indicated by a † (Kruskal-Wallis with post-hoc Mann-Whitney U; *H* = 6.4697, *p* < 0.05, with mean ranks of 30, 23.7143, and 17.6923 for Normal, Mild/Moderate CI and Severe CI respectively.)

| | Normal ($n = 31$) | Mild/Moderate CI ($n = 7$) | Severe CI ($n = 13$) |
| --- | --- | --- | --- |
| Age in months (SD) | 36.18 (19.87)<sup>†</sup> | 26.76 (17.06) | 20.37 (18.56)<sup>†</sup> |
| Male:Female Ratio | 20:11 | 6:1 | 6:7 |
| Ethnicity |  |  |  |
| Asian | 2 (6%) | — | 1 (8%) |
| Black | — | 1 (14%) | — |
| White (U.K./European) | 29 (94%) | 6 (86%) | 12 (92%) |
| Antiepileptic Drugs |  |  |  |
| None | 3 (10%) | 1 (14%) | — |
| Monotherapy | 26 (84%) | 6 (86%) | 9 (69%) |
| Polytherapy | 2 (6%) | — | 4 (31%) |
| Focal Seizures | 12 (39%) | 3 (43%) | 4 (31%) |
| Generalized Seizures | 18 (58%) | 2 (28%) | 9 (69%) |
| Generalized and Focal | 1 (3%) | 2 (28.5%) | — |
| Epilepsy aetiology |  |  |  |
| Cryptogenic | 3 (10%) | 1 (14%) | 5 (38%) |
| Idiopathic | 24 (77%) | 4 (57%) | 1 (8%) |
| Symptomatic | 3 (10%) | 2 (29%) | 7 (54%) |
| Unknown | 1 (3%) | — | — |
| Cognitive $z$-score (SD) | -0.05 (0.66) | -1.41 (0.20) | -2.9 (0.27) |

A retrospective analysis was done on 32-channel, unipolar montage with average reference captured routine EEGs. EEGs were recorded at 20 scalp electrodes (FP1, FP2, FPz, F3, F4, F7, F8, Fz, C3, C4, Cz, P3, P4, Pz, T3, T4, T5, T6, 01, 02), eight auxiliary electrodes (AUX1-8), two grounding (A1, A2) and two ocular electrodes(PG1, PG2).

### 2.2. Pre-processing

EEG recordings were pre-processed in MATLAB using the Fieldtrip toolbox [17]. The EEG had a sampling rate of approximately 511 Hz. Recordings were re-referenced to a common average reference (CAR), and bandpass filtered between 0.5-45 Hz in Fieldtrip. The resting-state data was split into non-overlapping, two second long sub-trials; long enough to pick up any resting-state network activity, while still fitting at least one full period of the lowest included frequency.

Prior to data processing, seizure activity in the EEGs were confirmed by clinicians. Whole trials which contained seizure activity were excluded from the analysis, rather than excluding only sections of trials with evident seizure activity. This helped guarantee that all network trials were derived from a minimum of two continuous seconds of seizure-free EEG. The small time window helped to balance removing large amounts of useful EEG data, while retaining enough data to characterize the frequencies present.

Standard EEG artefacts were rejected using a 2-step approach with manual and automatic rejection. Manual artefact rejection first removed clear outliers in both trial and channel data based upon high variance values (*var* > 10^6^). Muscle, jump and ocular artefacts were then automatically identified using strict rejection criteria relative to the Fieldtrip default suggested values [17] (Fieldtrip release range R2015-R2016b, *z*-value rejection level *r* = 0.4). All trials containing EEG artefacts were excluded from analysis. For subjects, we averaged across all trials at each frequency band, to help reduce potential bias and variance resulting from our selection of a shorter analysis window.

A narrow band (2-Hz wide) approach was used in analysis of clean EEG data, similar to work done by Miskovic et al. [18]. Segmenting the frequency range into these narrow bands (e.g. 1-3 Hz, 3-5 Hz,…) provided a data-driven approach to interrogate networks across subjects. The a priori nature of the investigation avoided attempts at equivocating the (likely heterogeneous) impact of epilepsy, development, medication etc. on each child’s spectral EEG composition. While such narrow bands may eschew some physiological interpretations by not adhering to classical frequency bands, the narrow bands promoted identification of mainly robust, common network abnormalities across the heterogeneous CWEOE population. If significant network abnormalities were identified in these narrow frequency bands (after correction for multiple comparisons, age and spurious correlations) then the identified results were likely a strong effect.

### 2.3. Network Coupling Analysis

The processed data was analyzed using functional EEG graph analysis, based on ‘functional links’ connecting any pair of EEG channels *i* and *j*, derived from the cross-spectrum of the data. Appendix A provides the detailed, formal definitions for the cross-spectrum and the network analysis methods described below. A summary of these definitions are included here for clarity. In brief, this study selected several measures of dependencies in EEG recordings, created graph networks based on these measures and characterized the created networks to identify candidate biomarkers for classification and identification of CI in CWEOE.

This study investigates three connectivity analysis methods building from the cross-spectrum viz: (1) the imaginary part of coherency (ICOH) [19], (2) phase-slope index (PSI) [20], and (3) weighted phase-lag index [21, 22].

ICOH is a standard measure in functional network analysis [19]. ICOH is well documented, and has been shown to provide direct measures of true brain interactions from EEG while eliminating self-interaction and volume conduction effects [19]. A weakness of ICOH, however, is its dependence on phase-delays, resulting in identifying functional connections only at specific phase differences between signals, while completely failing for others [21, 22, 23].

The PSI [20] was selected as a complementary alternative to ICOH for analysis. In practice, the PSI examines causal relations (temporal order) between two sources for signals of interest, e.g. *s_i_* and *s_j_* [20]. PSI exploits the phase differences between the sources to identify the ‘driving’ versus ‘receiving’ relationship between the sources [20]. Their average phase-slope differences are used to identify functional links [20]. Importantly, unlike ICOH, the PSI is equally sensitive to all phase differences from cross-spectral data [20]. However, the PSI equally weights contributions from all phase differences, meaning even small phasic perturbations are equal to the (defining) large perturbations.

Therefore the weighted phase-lag index (WPLI) was included as a third comparative measurement for analysis [21, 22]. The standard phase-lag index (PLI) [21] is a robust measure derived from the asymmetry of instantaneous phase differences between two signals, resulting in a measure which is less sensitive to volume conduction effects and independent of signal amplitudes [21]. The PLI ranges between 0 and 1, where PLI of zero indicates no coupling (or coupling with a specific phase difference; see [21] for details), while a PLI of 1 indicates perfect phase locking [21]. The PLI’s sensitivity to noise, however, is hindered as small perturbations can turn phase lags into leads and vice versa [22].

A weighted version of the PLI was introduced (weighted PLI; WPLI) [22] to counter this effect. The WPLI adds proportional weighting based on the imaginary component of the cross-spectrum [22]. The proportional weighting alleviates the noise sensitivity in PLI. The WPLI, like the PSI, helps capture potential phase-sensitive connections present in EEG networks from another perspective.

### 2.4. Adjacency Matrices and Sub-Networks

The estimated functional connectivity between channel pairs *i* and *j* comprising the weighted functional network of a subject can be represented by an adjacency matrix. The functional connections found for the ICOH, PSI, and WPLI measures were therefore represented via adjacency matrices in the analysis below. A set of adjacency matrices for a representative normal and impaired cognition child in the range of 5-9 Hz are included in Apppendix B, Figures B.5 and B.6, respectively.

Methodological choices associated with constructing and comparing graphs from the adjacency matrix can introduce bias in the network analysis (see [24, 25, 26] for details). Therefore, we used two methods for defining unbiased sub-networks of the functional EEG for comparison and analysis: the Minimum Spanning Tree (MST) [24] and the Cluster-Span Threshold (CST) [27].

The MST is an acyclic, sub-network graph which connects all nodes (electrodes) of a graph while minimizing link weights (connectivity strength) based on applying Kruskal’s algorithm on the weighted network [24, 28]. In brief, the algorithm orders the link weights in a descending manner (i.e. from strongest connection to weakest), constructing the MST by starting with the largest link weight and adding the next largest link weight until all nodes, *N*, are connected in an acyclic sub-network with a fixed density of *M* = *N* – 1 [24]. After construction of the sub-network, all weights are assigned a value of one [24]. In this manner, the MST is able to efficiently capture a majority of essential properties underlying a complex network in an unbiased sub-network [24].

Exploiting the properties of the MST is a standard technique common in recent publications exploring brain networks [24]. However, since the MST naturally leads to sparse networks in the data due to its acyclic nature, and that in some occasions more dense networks may be preferable, there is potentially real brain network information lost in the MST based EEG graph analysis [29].

By contrast, the CST creates a similar sub-network, but balances the proportion of cyclic ‘clustering’ (connected) and acyclic ‘spanning’ (unconnected) structures within a graph (for details see [27]). This balance thus retains naturally occurring ‘loops’ which can reflect dense networks without potential information loss [29] while still producing an unbiased sub-network for analysis. Figure 2 illustrates a topographical example of EEG channels connected via MST and CST networks for a randomly selected child. Differences in sparsity between the acyclic MST and the cyclic CST sub-networks can readily be seen in Figure 2. Both the MST and CST are binary sub-networks, which have additional advantages over weighted networks, e.g. the adjacency matrix [24, 27, 29].

**Figure 2:**
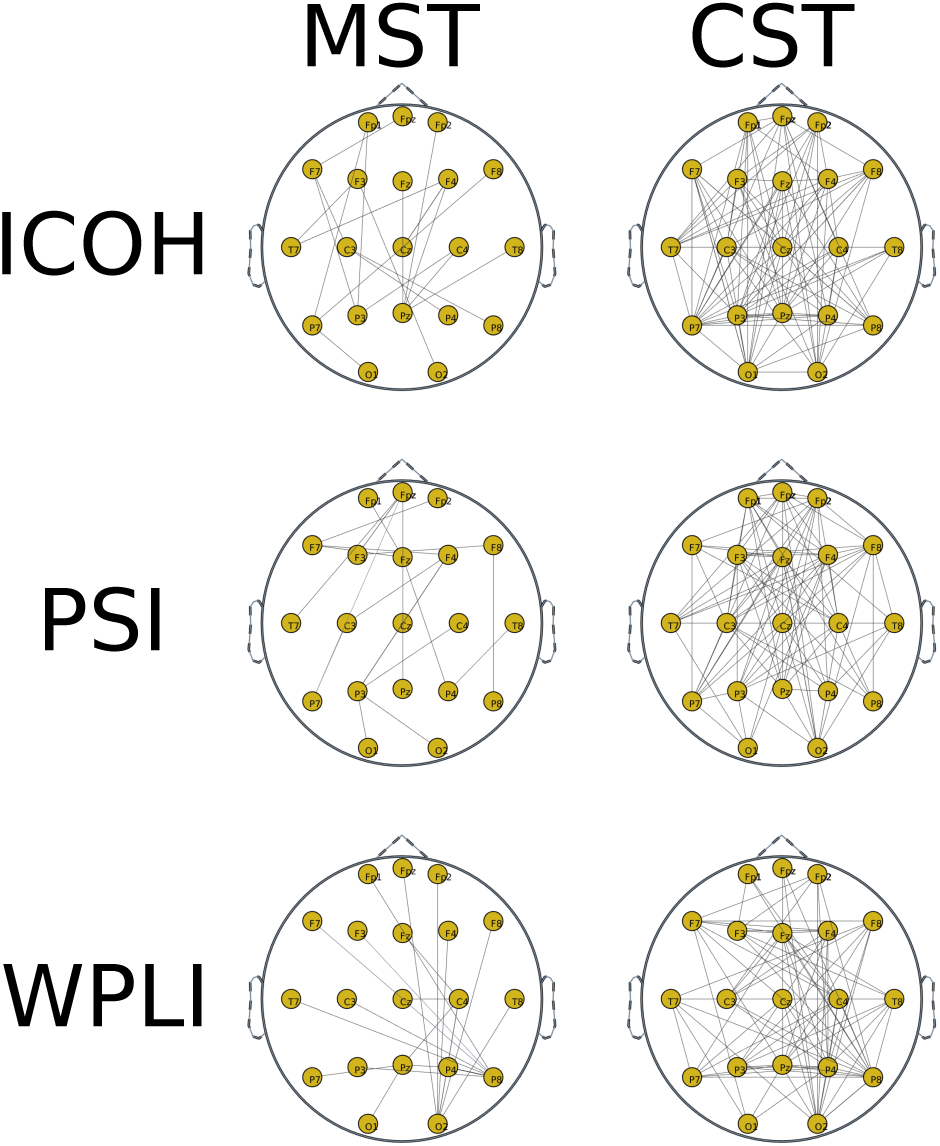
Illustrative examples of the MST and CST sub-network graphs of ICOH, PSI and WPLI for a randomly selected child. EEG channels are displayed as nodes, with functional connections displayed for each combination of sub-network and connectivity measure.

For each combination of sub-networks and connectivity definitions above (e.g. MST-ICOH, CST-ICOH, MST-PSI, etc.) four network metrics were investigated for correlation to the cognition standard score measures. To help reduce potential selection bias, network metrics for analysis were agreed upon a priori. Metrics were chosen to account for distinct network properties (e.g. the shape of the network, the critical connection points in the network etc.) with (relatively) little inter-correlation. Due to the natural exclusion/inclusion of cycles, the network metrics differ for the MST and CST, respectively. However, all metrics across sub-networks were selected to be comparable regarding network properties. Pictorial examples of the selected network metrics, alongside short definitions, are outlined in Figure 3.

**Figure 3:**
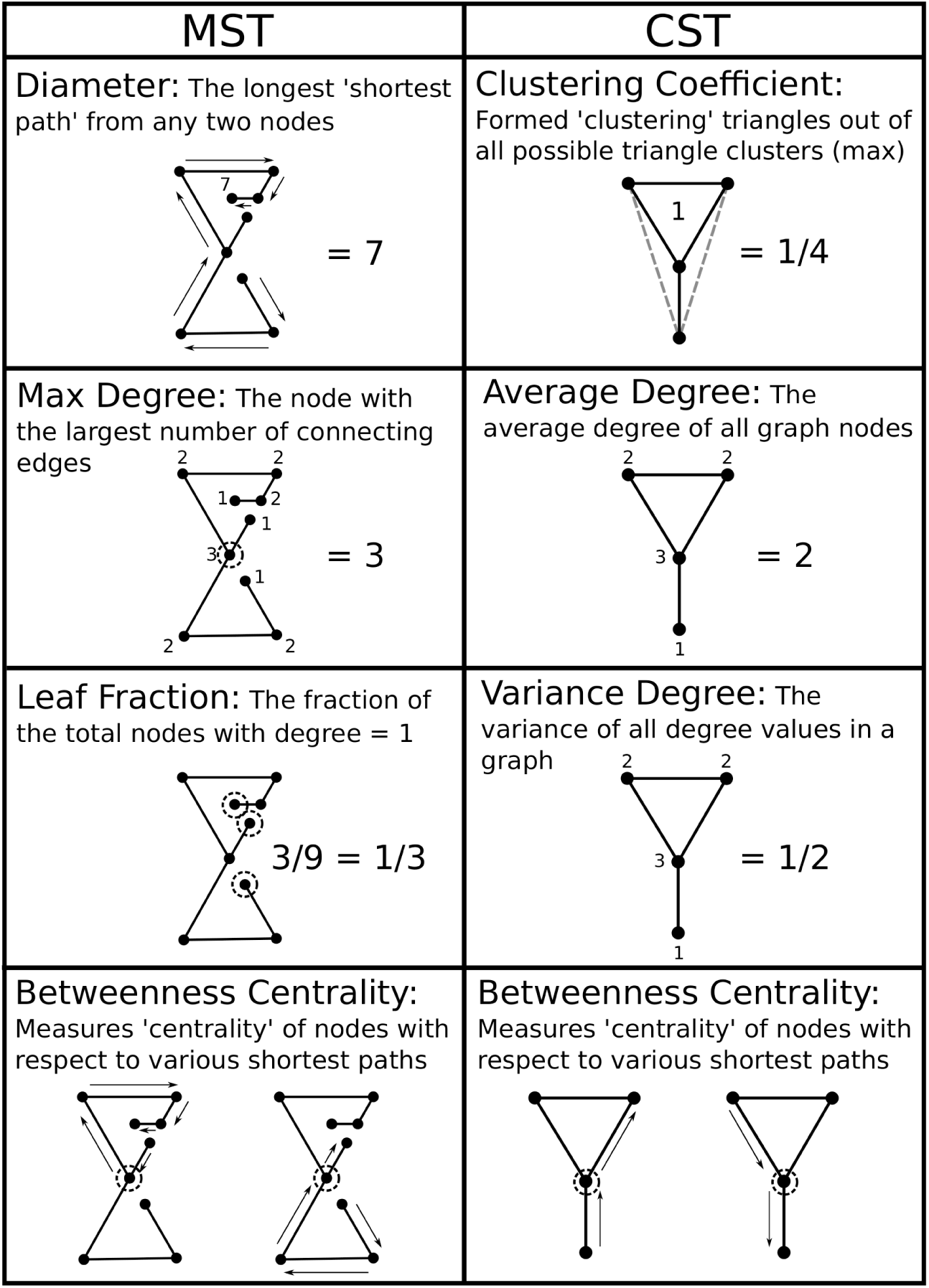
Illustration of all graph analysis metrics for the Minimum Spanning Tree (MST) and Cluster-Span Threshold (CST) networks using simple example graphs. Nodes (dots) represent EEG channel electrodes. Edges (lines) represent functional interactions between EEG channels identified by a connectivity measure, e.g. ICOH/PSI/WPLI.

### 2.5. Statistical Analysis

Statistical analysis was done using Matlab 2015a. Correlation between individual network metrics and the cognition standard score was measured using Kendall’s tau (*τ*) [30]. Kendall’s *τ* calculates the difference between concordant and discordant pairs[30, 31], and is ideal for describing ordinal or ranking properties, like the normalized cognition standard score. Its design is also relatively robust to false positive correlations from data outliers [30, 31], providing additional mitigation to spurious correlations in the results. Furthermore, as Kendall’s *τ* is a non-parametric hypothesis test it did not rely on any underlying assumptions about the distribution of the data. Therefore our correlation analysis was robust to any potential ceiling, floor or skewed distribution effects present in the reported cognition standard score measures.

Correlation trends are reported both as uncorrected *p* < 0.05 values, and with multiple comparison (Bonferroni) corrections, similar in style to previous literature [32]. For each frequency bin (2-Hz wide) and network, we compared and corrected for the 4 separate graph measures using the Bonferroni technique (i.e. we set *p* = 0.05/4 = 0.0125 as the threshold for significance). Dependency was assumed across the small 2-Hz frequency bins, similar in principle to [32], and as such we do not include the frequency bins in the Bonferroni correction. Correlations which are found to be potentially significant under this assumption are indicated by the † symbol for Bonferroni corrections.

### 2.6. Classification

A multi-class classification scheme was devised using the Weka toolbox [33, 34]. Class labels of *normal*, *mild/moderate CI*, and *severe CI* were applied.

Primary feature selection included all correlations identified by the statistical analysis, thereby allowing potential interpretation of the retained network features. Then, a second feature selection phase using nested 5-fold cross-validation selected prominent features via bi-directional subspace evaluation [35]. Within this nested cross-validation, features identified as important in > 70% of the folds were selected for use in classification.

Due to natural skew of the data (towards normalcy), and the context of the classification problem (e.g. misclassifying different classes has various implications), a cost-sensitive classifier was developed [36]. In order to properly develop such a classifier, an appropriate cost matrix needed to be identified.

Using guidelines outlined in literature [36], the cost matrix in Table 2 was developed, with predicted classes on the rows and real classes on the columns.

**Table 2:**
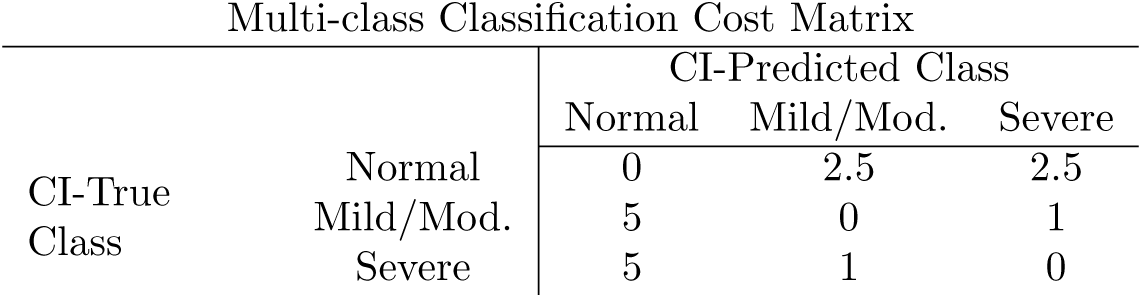
Weighted cost matrix for misclassification of cognitive impairment (CI) for normal (±1 SD), mild/moderate (–1 to –2 SD) and severe (< –2 SD) classes. Rows represent true class labels, with columns as the predicted classification labels.

The defined matrix satisfies several key concerns in multi-class cost-matrix development [36]. The weights on misclassification were carefully selected to reflect probable clinical concerns in classification with guidance from paediatric neurologists (RC, JS). The cost for incorrectly classifying an impaired child as normal was twice as heavy compared to misclassifying a normal child into either impaired group, which was still significantly more punishing than correctly identifying impairment and only misclassifying between mild/moderate or severe impairments. These weighted values prioritized correctly including as many ‘true positive’ CWEOE with CI, i.e. increasing sensitivity, followed by a secondary prioritization upon being able to discern the level of CI. These boundaries provide a more clinically relevant classification context in the analysis.

Using the selected features and developed cost-sensitive matrix, a nested 5-fold cross-validation trained a simple *K*-Nearest Neighbour (KNN) classifier, with *N* = 3 neighbours and Euclidean distance to minimize the above costs. By demonstrating our proof-of-concept results with a simple classifier first, e.g. KNN, we aimed to highlight that network response found from our analysis pipeline was likely robust. A repeated ‘bagging’ (Boostrap Aggregation [37]) approach was used to reduce variance in the classifier at a rate of 100 iterations/fold. Results were evaluated upon their overall classification accuracy and total penalty costs (e.g. sum of all mistakes based on the cost matrix).

Random classification and naive classification (e.g. only choosing a single class for all subjects) were included for comparison. In this study, random classification refers to classification of any ‘true’ class label to a randomly selected ‘predicted’ class label. Based on the distribution of subjects into the classes, a ’chance’ level for each class is used to assign the ’predicted’ label at random. Naive classification (e.g. single-class classification), assumes that all subjects belong to only one class. Classification accuracy and misclassification penalties are then calculated based on the presumed (single) class assignment. This study looked at naive classification for each class label, and have reported comparisons to each possible naive classification.

## 3. Results

Of 64 children enrolled into the parent study, 13 were excluded from the current study due to corrupted EEG data and inconsistent or incompatible EEG acquisition parameters. There were data available for analysis on 51 children (32:19 male-to-female ratio, mean age and SD of 30.85 ± 20.08 months). On average approximately 455 ± 325 two second trials were used for each child in the analysis, totalling 15.16 ± 11.87 minutes of resting-state EEG data for each child. Thirty-one children had normal cognition, 7 had mild/moderate CI, and 13 had severe CI.

### 3.1. Correlation Analysis

Each combination of functional link analysis (ICOH/PSI/WPLI) and sub-network selection (MST/CST) techniques uncovered likely correlations between at least one network metric (outlined in Figure 3) and the cognition standard score measures. A summary of the significant correlations between the MST metrics and the standard scores are shown in Table 3. All MST correlations were in the medium to high frequency range, 9 – 31 Hz, with no significant results in lower frequencies. Activity above approximately 9 Hz is outside of the expected range for the delta, theta and alpha bands in young children [38, 39]. Sets of contiguous frequency bands with significant correlations were found in the ICOH and PSI connectivity measures, and are reported together as a single frequency range. Overlapping correlations retained at significant levels after partial correlation correcting for age are also reported for the MST using a modified Kendall’s *τ*.

**Table 3:** Summary of Kendall’s *τ* correlation trends between various graph metrics and the cognition standard scores using the Minimum Spanning Tree (MST). For all values |*τ*| was between 0.201 and 0.310; mean = 0.239 ± 0.0278 and uncorrected *p* < 0.05. Significant values across contiguous narrow-band frequencies have been grouped together for ease of interpretation. ^†^Significant with Bonferroni correction at the level of frequencies. ^∗^ Significant after partial correlation correction to age of subjects, via modified *τ* with uncor-rected *p* < 0:05.

| MST analysis of cognition standard score measures |  |  |  |
| --- | --- | --- | --- |
| Network Type | Network Measurement | Frequency Range(s) (Hz) | Correlation ($\bar{\tau} \pm SD$) |
| ICOH | Diameter | — | — |
| ICOH | Maximum Degree | — | — |
| ICOH | Leaf Fraction | — | — |
| ICOH | Betweenness Centrality | 13-17 Hz | $-0.231 \pm 0.001$ |
| PSI | Diameter | 9-19 Hz | $0.239 \pm 0.032^{\dagger*}$ |
| PSI | Maximum Degree | 11-13 Hz | $-0.232 \pm 0.000^*$ |
| PSI | Maximum Degree | 15-17 Hz | $-0.258 \pm 0.000^{\dagger*}$ |
| PSI | Maximum Degree | 21-23 Hz | $-0.219 \pm 0.000$ |
| PSI | Leaf Fraction | 11-13 Hz | $-0.201 \pm 0.000$ |
| PSI | Leaf Fraction | 15-19 Hz | $-0.246 \pm 0.003$ |
| PSI | Betweenness Centrality | 9-13 Hz | $-0.218 \pm 0.012^*$ |
| PSI | Betweenness Centrality | 17-19 Hz | $-0.259 \pm 0.000^{\dagger*}$ |
| WPLI | Diameter | — | — |
| WPLI | Maximum Degree | 29-31 Hz | $-0.310 \pm 0.000^{\dagger*}$ |
| WPLI | Leaf Fraction | — | — |
| WPLI | Betweenness Centrality | 23-25 Hz | $0.223 \pm 0.000$ |

Similarly, significant correlations between the CST metrics and the cognition standard scores are shown in Table 4. Several significant CST metrics exist in the lower frequency range (< 9 Hz), indicating a potential sensitivity of the CST to lower frequencies. No sets of continuous frequency bands were discovered, but several sets were trending towards this phenomenon within ICOH. Multiple overlapping correlations remaining after partial correlation correction for age from the modified *τ* in the CST at lower frequencies indicate additional sensitivity.

Both the MST and CST demonstrate high sensitivity in the phase-dependent measures (PSI, WPLI) compared to the standard ICOH.

### 3.2. KNN Classification

Based upon CST’s sensitivity, a preliminary classification scheme assessed the potential predictive qualities of the CST network metrics in identifying CI classes. The relative quality of the classifications are examined using classification accuracy and total ‘cost’ (i.e. penalty for misidentification) [36].

The subset of CST metrics for classification, identified from significant correlations and chosen via cross-validated feature selection, included five network metrics across the three connectivity measures. For ICOH, the identified subset selected was the betweenness centrality at ranges 11-13 and 19-21 Hz alongside the clustering coefficient at a range of 15-17 Hz. The subset also included the PSI average degree at 13-15 Hz and the WPLI variance degree from 1-3 Hz. These results indicate specifically which network metrics, from a machine-learning perspective, contributed the most information for building an accurate classification model. As such, the classifier was trained specifically, and only, using these 5 key metrics. An illustrative example of these 5 selected network metrics (e.g. features) are shown in Figure 4 as scatter plots. When training the classifier, these network features are used to identify the underlying patterns not readily observed, and are incorporated into guiding the machine learning algorithm.

**Figure 4:**
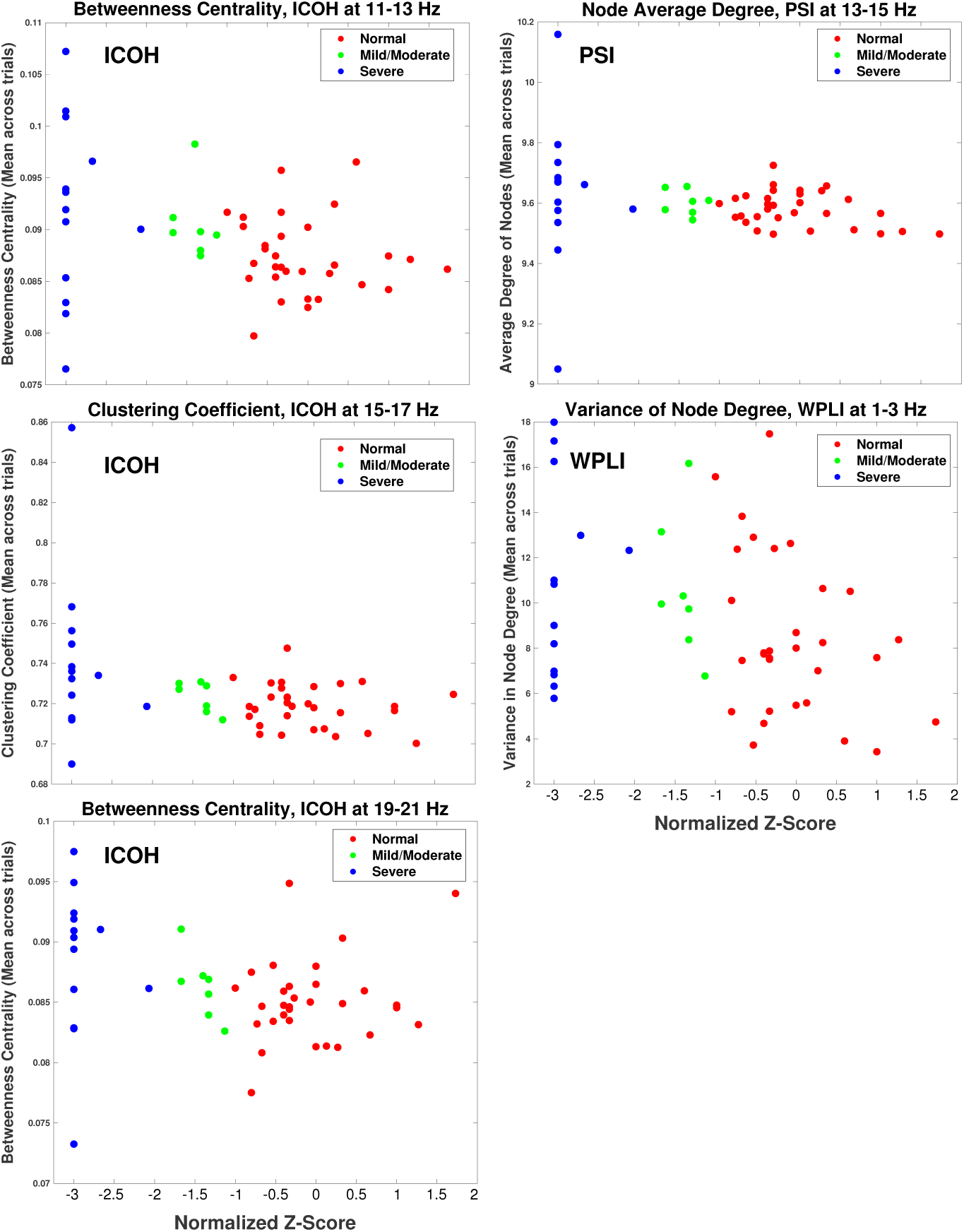
Scatter plot displaying the distribution of children for each of the 5 features used in training the KNN classification. Each panel displays network values on the y-axis, with the normalized cognition standard score (*z*-score) on the x-axis. Children classified into normal, mild/moderate CI and severe CI classes are displayed in red, green and blue respectively.

It bears repeating that Kendall’s *τ* is a non-parametric significance test, which means it does not rely on an underlying assumption of a specific type of distribution in the data. Therefore, Kendall’s *τ* correlation was robust to the apparent flooring effect seen in the severe CI class, as it utilizes concordant and discordant pairs. Therefore our choice of features from the statistical analysis remains unaffected.

The resulting confusion matrix from the 5-fold cross-validated, cost-sensitive classification analysis is seen in Table 5, with key summary

The overall classification accuracy was defined as the number of true label classes correctly predicted by the classifier, e.g. the true positive diagonal of Table 5. Presently, approximately 36 of the 51 children’s cognitive class (e.g. normal, mild/moderate CI, severe CI) were correctly predicted, giving a total accuracy of the classifier at 70.6%. Using Table 2, an overall ‘cost-penalty’ value was calculated at 38 points, based on the children who were misclassified, i.e. their cognitive class was not correctly predicted.

**Table 4:** Summary of Kendall’s *τ* correlation trends between various graph metrics and the cognition standard scores using the Cluster-Span Threshold (CST).For all values |*τ*| was between 0.201 and 0.290; mean = 0.237 ± 0.033, and uncorrected *p* < 0.05. Significant values across contiguous narrow-band frequencies have been grouped together for ease of interpretation. ^†^Significant with Bonferroni correction at the level of frequencies. ^∗^ Significant after partial correlation correction to age of subjects, via modified *τ* with uncor-rected *p* < 0:05.

| CST analysis of cognition standard score measures |  |  |  |
| --- | --- | --- | --- |
| Network Type | Network Measurement | Frequency Range(s) (Hz) | Correlation ($\bar{\tau} \pm SD$) |
| ICOH | Clustering Coefficient | 15-17 Hz | $-0.290 \pm 0.000^{\dagger*}$ |
| ICOH | Average Degree | — | — |
| ICOH | Variance of Degree | 13-15 Hz | $-0.200 \pm 0.000$ |
| ICOH | Variance of Degree | 21-23 Hz | $-0.203 \pm 0.000$ |
| ICOH | Betweenness Centrality | 11-13 Hz | $-0.273 \pm 0.000^{\dagger*}$ |
| ICOH | Betweenness Centrality | 15-17 Hz | $-0.241 \pm 0.000$ |
| ICOH | Betweenness Centrality | 19-21 Hz | $-0.203 \pm 0.000$ |
| PSI | Clustering Coefficient | — | — |
| PSI | Average Degree | 13-15 Hz | $-0.210 \pm 0.000$ |
| PSI | Variance of Degree | 15-17 Hz | $-0.277 \pm 0.000^{\dagger*}$ |
| PSI | Variance of Degree | 21-23 Hz | $-0.217 \pm 0.000$ |
| PSI | Betweenness Centrality | 5-7 Hz | $0.204 \pm 0.000^*$ |
| PSI | Betweenness Centrality | 15-17 Hz | $-0.248 \pm 0.000$ |
| WPLI | Clustering Coefficient | 1-3 Hz | $-0.236 \pm 0.000^*$ |
| WPLI | Clustering Coefficient | 17-19 Hz | $0.287 \pm 0.000^{\dagger*}$ |
| WPLI | Average Degree | — | — |
| WPLI | Variance of Degree | 1-3 Hz | $-0.236 \pm 0.000^*$ |
| WPLI | Betweenness Centrality | — | — |

**Table 5:** Resulting confusion matrix from the 5-fold cross-validated, cost-sensitive classification scheme for all *n* = 51 children based on costs in Table 2. Rows represent true class labels, with columns as the predicted labels from the classification. Bold values along the diagonal show true positive classification results, where actual and predicted cognitive classes were accurately identified. Italicized values indicate children predicted to have CI, i.e. mild/moderate or severe class, by the classification scheme.

| Confusion Matrix from Classification Results |  |  |  |  |
| --- | --- | --- | --- | --- |
|  |  | CI-Predicted Class |  |  |
|  |  | Normal | Mild/Mod. | Severe |
| CI-True Class | Normal | **26** | *2* | *3* |
|  | Mild/Mod. | **2** | **3** | *2* |
|  | Severe | **1** | *5* | **7** |

**Table 6:**
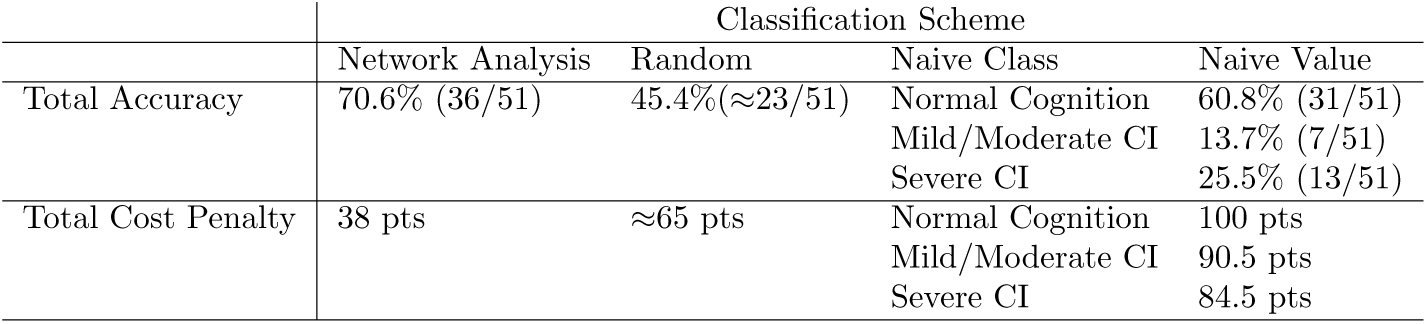
Summary table of overall classification accuracies and total cost penalty for the proposed network analysis, random classification, and naive (single class) classification. Naive classification is split to show overall classification accuracy and cost penalties if all children were assigned as normal cognition, mild/moderate CI or severe CI classes. Total accuracy includes the approximate number of children with true positive predictions, out of total number of children evaluated.

The expected random classification accuracy is based on the distribution of individuals belonging to each class, i.e. 31, 7 and 13 children for the *normal*, *mild/moderate* and *severe* classes respectively. Random accuracy would be expected at 45.4%, with cost-penalty varying depending on misclassification distributions. Using the average misclassification penalty and the percentage of misidentified children (approximately 28 of the 51 subjects), the cost-penalty would be at least 65 points.

Naive, or one-class classification assumes all subjects belong to a single class only. For example, if all children were considered to only belong to the ‘normal’ cognition class (i.e. naively classified as normal), then exactly 31 of the 51 children (those whose true class is ‘normal’-the first row of Table 5) would be correctly identified, giving a naive classification accuracy of 60.8%. Repeating this naive classification scheme for mild/moderate and severe classes provides naive classification accuracies of 13.7% (7/51), and 25.5% (13/51) respectively. Similarly, the total cost-penalty for each naive classification would be 100, 90.5 and 84.5 points respectively, using the same procedure and the penalty costs from Table 2.

Overall, the results indicate gains in classification accuracy and a reduced total penalty as compared to both random and naive classification. This is summarized in Table 6.

## 4. Discussion

The main finding of this study is the development of novel methods towards identifying a potential computational biomarker for CI in CWEOE. The automated and quantitative nature of the processing chain, ability to appropriately predict CI classes, and its use of routinely acquired EEG data make the proposed methods an attractive proposition for clinical applications. Our results indicate a substantial pool of potential characteristics might be identified using the proposed methods with several network analysis and filtering combinations.

The breadth of these combinations emphasizes the general suitability of EEG networks in identifying possible CI markers in CWEOE.

Flexibility in sensitivity and robustness of particular networks to features of interest is an advantage of this analysis. For instance, the sensitivity of phase-dependent connectivity measures, e.g. PSI and WPLI, was more prevalent compared to standard ICOH. This is not surprising as phase-oriented measures were developed to improve upon phase ambiguities in traditional ICOH measurements [20, 23]. In addition, the sensitivity of PSI in picking up significant correlations can be attributed in part to its equal treatment of small phase differences in leading and lagging signals [20]. Such small phase differences contribute equally in PSI, while counting for proportionally less in the WPLI by definition [22, 21]. By construction, the WPLI results are substantially more robust to noise and small perturbations in phase, through proportionally reflecting phase differences in network connections with appropriate weights, providing results for only large phase differences. Together these measures reflect trade-off choices between sensitivity and robustness for network analysis.

Of interest for paediatric populations is the CST’s capability to identify low frequency correlations in phase-dependent coherency measures. Both the PSI and WPLI demonstrate sensitivity to lower frequencies, not present in the ICOH or MST in general. This is critical considering that in preschool children lower frequencies typically contain the bands of interest present in adult EEGs, e.g. the delta/theta/alpha bands [38, 39]. During development these bands shift to higher frequencies [40], reflecting a large scale reorganization of the endogenous brain electric fields and suggesting a transition to more functionally integrated and coordinated neuronal activity [18]. The (low) chance of all such significant findings being spurious is of less detriment than the potential loss of impact for disregarding the findings if at least one of them is true. The sensitivity to detect network disruptions already present in these critical bands in CWEOE provide high value in adjusting potential therapeutic and treatment strategies for clinicians.

The identified subset of metrics for classification provide additional information. All of the features in the subset reflected distribution measures of hub-like network structures in the brain, relating to the balance between heterogeneity and centrality within the network. The implicated metrics, other than the variance degree, corresponded to measures identifying local, centralized ‘critical’ nodes in a network. Their negative correlation to the cognition standard scores imply that children with more locally centralized brain networks, and consequently with less well distributed hub-like structures, are more likely to have corresponding cognitive impairment. This is reasonable, since if there exists a small set of central, critical hubs responsible for communication across the brain, disruption of these critical points (e.g. due to seizure activity) would have severely negative effects on communication connections. This is also supported by the negative correlation in the variance degree metric in the WPLI. The variance degree can be interpreted as a measure of a network’s heterogeneity [41]. As such, the negative variance degree in the low (1-3 Hz) frequency range may reflect stunted cognitive development, as normal maturation is associated with reduced activation in low frequencies [42, 38, 43, 39, 44], implying a decrease in local connectivity and heterogeneity of the networks. This compliments the above conclusions, suggesting a sensitivity in the likely well-centralized networks to significant disruptions by epilepsy. The disrupted networks may then be reflected by the continued heterogeneity and local connectivity of low frequency structures in impaired children.

Being able to predict the likely extent of CI using the identified markers could provide an advantageous tool for clinicians. Specifically, being able to pair specific network features to an effective prediction of CI would allow clinicians to retain the interpretability of the chosen network features while providing a tool to quickly and objectively separate similar cases. To this end, the cost-sensitive, simple KNN classifier explored in this work illustrates an early step towards this aim. Evaluating the network-based classifier results show the analysis was successful at two levels. First, the proposed classifier was able to generally identify cognitively normal children from impaired children, when grouping the mild/moderate CI and severe CI classes. This is seen in the first column of Table 5 where only three impaired children are misidentified as ‘normal cognition’, giving a sensitivity of 85%. In other words, 17 of the 20 actual impaired children were correctly identified as belonging to either the mild/moderate or severe CI classes, demonstrating that the proposed network analysis and classifier was largely successful with respect to predicting children with some form of impaired cognition, based on using the standard score definition. Similarly, only five normal children were misidentified as generally impaired (i.e. classified to either the mild/moderate or severe CI classes; top row of Table 5), giving a specificity of approximately 84% (26/31) for appropriately identifying children in the range of normal cognition. In addition, the network coupled classifier was able to separate out cases of mild/moderate impairment from severe impairment decently, with > 50% of impaired children correctly predicted. Thus, the proposed classifier and associated methods provide considerable sensitivity (85%) and specificity (84%) for clinicians in determining potential CI, while still remaining relatively accurate in separating CI according to severity.

Statistical analysis in this manuscript was utilized as a first-pass means to reduce the potential feature space for classification. Through identifying potentially significant networks of interest, the number of features to test in the classification step was substantially reduced. Through the statistical filter, we were able to select pertinent features from a relevant and manageable feature space. Future endeavours could refine such features, based on different choices for the statistical analysis. Using a more rigid/flexible analysis could lead to further culling/relaxation of the feature space and provide an adjustable framework for examining network property changes in CWEOE. Other future work could include alternative narrow-band frequency binning and less strict automated rejection methods. Significant correlations across sets of consecutive (and nearly consecutive) frequency bands indicate likely targets for potential followup studies. Further development of a more complex classification scheme could help improve the second tier discrimination of the proposed classifier, at the level of discerning between the cognitive impairment types (e.g. mild/moderate CI from severe CI). A thorough investigation into incorporating and comparing additional classifiers is also a potential avenue for expansion of this research.

The NEUROPROFILE cohort was advantageous in that formal neuropsychological testing was coupled with EEG recordings, making it ideal for this investigation. However, there are study limitations. Although this novel study used routine clinical EEGs used in the diagnosis of incidence cases of CWEOE, the three classes of normal, mild/moderate and severe impairment were unbalanced; this occurred naturally. The majority of the sample was taken from a population-based cohort, and mitigating potential influences from imbalanced data was taken into account as much as possible when conducting the research, e.g. through cost-sensitive analysis. Imbalanced data is not uncommon, but the unbalanced distribution of CI in the current study reflects findings in a true population-based cohort [45]. Furthermore, trialling this methodology in older children with epilepsy may be an avenue for future studies, to provide further insights as to the relationship between aetiology and CI, as well as provide additional replications of the proposed techniques.

## 5. Limitations

Within the studied cohort of CWEOE, the epilepsy type and aetiologies were heterogenous. Thus we are unable to determine if the model and methods used have greater or lesser predictive value in specific subsets. Testing in a larger, more homogeneous sample would provide clarification.

A gender disparity was noted within the normal cognition and mild/moderate CI groups. Although this study reflects a true population, further studies are needed to investigate this phenomena.

Note that the spectral components in the very low frequency narrow band (e.g.1-3 Hz) may not be fully reliable due to the small epoch length, i.e. two seconds. Information gained from the very low frequency band needs to be interpreted with some care, as spurious connections are more likely to be present. Again, however, the large number of trial epochs averaged for each child helped mitigate these potential spurious connections.

We recognize a limitation in our assumption of dependency between the frequency bins. While there is likely a strong local family dependency between the narrow bins, the endpoints on our frequency spectrum may not have as strong of a relation. Therefore, significance at these level should be considered carefully as they are more likely to be a false positive. However, the robust nature of *τ* and our choice of features from a machine-learning perspective help to moderate potential impacts from this assumption on our results.

The use of a data-driven, narrow band approach in our analysis had a tradeoff of not using patient-specific frequency ranges for each child. Future studies could be done to investigate how individualized frequencies, e.g. using individual alpha frequencies (IAF), could be aligned, interpreted and correlated when assessing network abnormalities in the CWEOE population.

## 6. Conclusions

This study introduced a novel processing chain based on network analysis for identifying markers of CI in CWEOE for the first time. Results from the study demonstrate these network markers in identifying critical structures of CWEOE with CI and illustrate their potential predictive abilities using preliminary classification techniques. Replication of the identified methods using other datasets, with alternative narrow-band frequency binning, less strict automated rejection methods, and including correlations with brain MRI abnormalities may bolster the generalizability and applicability of the proposed techniques.

## 7. Acknowledgements

The authors would like to thank the patients and families who participated in the NEUROPROFILES [45] study. Funding support for this project was provided by the RS McDonald Trust, Thomas Theodore Scott Ingram Memorial Fund, and the Muir Maxwell Trust.

## 8. Author Contributions

Javier Escudero and Richard FM Chin conceived of the presented ideas. Eli Kinney-Lang developed the theory, performed data analysis and interpretation, and designed the computational framework of the project under supervision of Richard FM Chin and Javier Escudero. Jay Shetty, Krishnaraya Kamath Tallur, Michael Yoong and Ailsa McLellan were involved in the methodology and collection of the original NEUROPROFILES dataset, including recruiting patients and requesting and reporting patient EEGs. Matthew Hunter was the lead author and investigator for the NEUROPROFILES project with senior supervision under Richard FM Chin. Eli Kinney-Lang wrote the manuscript and figures, with revision and comments provided by Matthew Hunter, Michael Yoong, Jay Shetty, Krishnaraya Kamath Tallur, Ailsa McLellan, Richard FM Chin and Javier Escudero. Final approval of this publication was provided by all authors.

## Appendix A. Network Coupling Definitions

Appendix A outlines the key network definitions and details for the presented analysis. For in-depth reviews see [46, 13], and for further reading [12, 47, 48].

### Cross-spectrum

Functional EEG connections are established through measures of interdependency between signals *s_i_* and *s_j_* [48] for any pair of EEG channels *i* and *j*. A common measurement for examining this interdependency is the cross-spectrum function *S_ij_* (*f*) [49, 19, 48]. Formally, let *x_i_*(*f*) and *x_j_* (*f*) be the complex Fourier transforms of the time series signals *s_i_* and *s_j_* for any pair (*i*,*j*) of EEG channels. Then the cross-spectrum can be calculated as

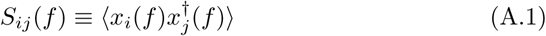

where † indicates the complex conjugation, and 〈〉 refers to the expectation value (also written as *E*{}) [19].

### Imaginary Part of Coherency (ICOH)

Coherency is defined as the normalized cross-spectrum[19]:

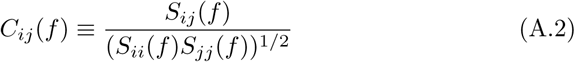

Therefore, the imaginary part of coherency is defined as [19]

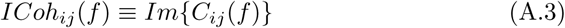

where *Im*{} refers to taking the imaginary part of the complex coherency measure.

### Phase-Slope Index (PSI)

The PSI is defined as:

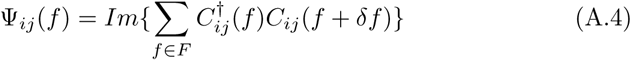

where *C_ij_*(*f*) is as defined in equation A.2, † indicates the complex conjugation, *δf* is the frequency resolution, and *f* ∈ *F* is the set of frequencies over which the phase-slope is calculated (see [20] for details).

### Phase-Lag Index

The PLI is defined as: [21, 22]

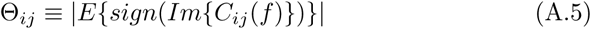

where *E*{} is the expectation, *sign* is the positive or negative sign, and *Im{C_ij_* (*f*)} is the same as ICOH (see equation A.3).

### Weighted Phase-Lag Index (WPLI)

The weighted PLI (WPLI) is defined as: [22]

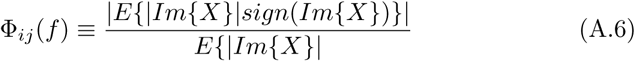

where *X* = *Im*{*C_ij_*(*f*)} = *ICok_ij_*(*f*).

## Appendix B. Supplementary Figures

**Figure B.5:**
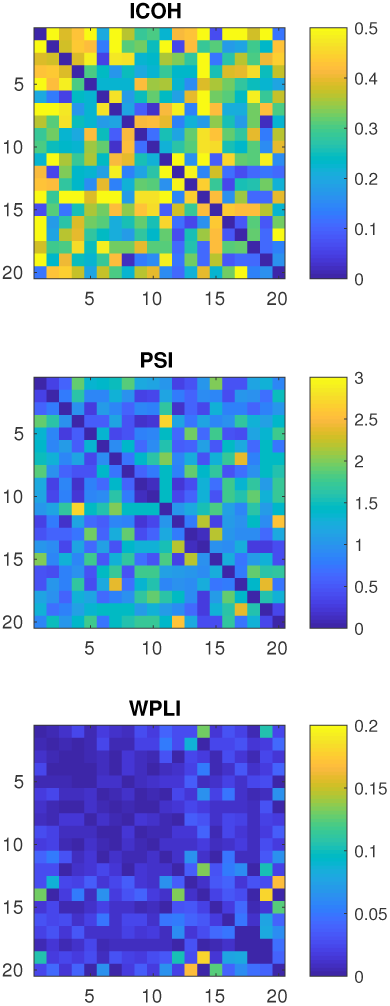
Adjacency matrices for a representative ‘normal cognition’ child calculated by ICOH, PSI and WPLI between the 5-9 Hz frequency range.

**Figure B.6:**
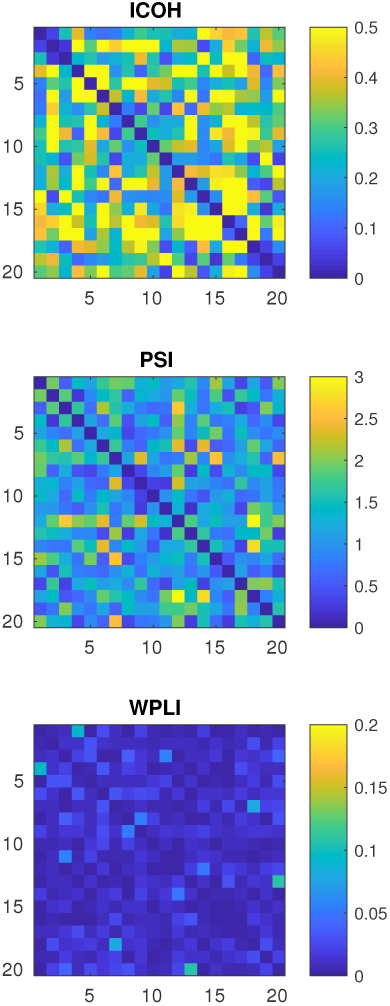
Adjacency matrices for a representative ‘impaired cognition’ child calculated by ICOH, PSI and WPLI between the 5-9 Hz frequency range.

